# Examination of the Shared Genetic Basis of Anorexia Nervosa and Obsessive-Compulsive Disorder

**DOI:** 10.1101/231076

**Authors:** Zeynep Yilmaz, Matthew Halvorsen, Julien Bryois, Dongmei Yu Eating Disorders Working Group of the Psychiatric Genomics Consortium, Tourette Syndrome/Obsessive-Compulsive Disorder Working Group of the Psychiatric Genomics Consortium, Laura M. Thornton, Stephanie Zerwas, Nadia Micali, Rainald Moessner, Christie L. Burton, Gwyneth Zai, Lauren Erdman, Martien J. Kas, Paul D. Arnold, Lea K. Davis, James A. Knowles, Gerome Breen, Jeremiah M. Scharf, Gerald Nestadt, Carol A. Mathews, Cynthia M. Bulik, Manuel Mattheisen, James J. Crowley

**Affiliations:** Department of Psychiatry, University of North Carolina at Chapel Hill, Chapel Hill, NC, USA; Department of Genetics, University of North Carolina at Chapel Hill, Chapel Hill, NC, USA; Department of Medical Epidemiology and Biostatistics, Karolinska Institutet, Stockholm, Sweden; Psychiatric and Neurodevelopmental Genetics Unit, Center for Genomic Medicine, Department of Psychiatry, Massachusetts General Hospital, Harvard Medical School, Boston, MA, USA; Department of Psychiatry, Faculty of Medicine, University of Geneva, Geneva, Switzerland; UCL Institute of Child Health, UCL, London, UK; Department of Psychiatry, Icahn School of Medicine at Mount Sinai, New York, NY, USA; Department of Psychiatry and Psychotherapy, University of Tuebingen, Tuebingen, Germany; Genetics & Genome Biology, The Hospital for Sick Children, Toronto, Ontario, Canada; Neurogenetics Section, Molecular Brain Science Department, Campbell Family Mental Health Research Institute, Centre for Addiction and Mental Health, Toronto, ON, Canada; Department of Psychiatry, University of Toronto, Toronto, ON, Canada; Groningen Institute for Evolutionary Life Sciences, University of Groningen, The Netherlands; Department of Translational Neuroscience, Brain Center Rudolf Magnus, University Medical Center Utrecht, Utrecht University, The Netherlands; Mathison Centre for Mental Health Research & Education, Cumming School of Medicine, University of Calgary, Calgary, Alberta, Canada; Vanderbilt Genetics Institute, Vanderbilt University Medical Center, Nashville, TN, USA; Division of Genetic Medicine, Department of Medicine, Vanderbilt University Medical Center, Nashville, TN, USA; Department of Psychiatry and Behavioral Sciences, Vanderbilt University Medical Center, Nashville, TN, USA; Department of Psychiatry and Zilkha Neurogenetic Institute, Keck School of Medicine, University of Southern California, Los Angeles, CA, USA; MRC Social, Genetic and Developmental Psychiatry Centre, Institute of Psychiatry, Psychology & Neuroscience, King’s College London, London, UK; Department of Psychiatry and Behavioral Science, Johns Hopkins University, Baltimore, MD, USA; Department of Psychiatry, Genetics Institute, University of Florida, Gainesville, FL, USA; Department of Biomedicine, Aarhus University, Aarhus, Denmark; The Lundbeck Foundation Initiative of Integrative Psychiatric Research (iPSYCH), Denmark; Department of Psychiatry, Psychosomatics and Psychotherapy, University of Würzburg, Würzburg, Germany; Department of Clinical Neuroscience, Karolinska Institutet, Stockholm, Sweden

**Keywords:** anorexia nervosa, obsessive-compulsive disorder, GWAS, genetic, SNP, heritability, meta-analysis

## Abstract

Anorexia nervosa (AN) and obsessive-compulsive disorder (OCD) are often comorbid and likely to share genetic risk factors. Hence, we examine their shared genetic background using a crossdisorder GWAS meta-analysis of 3,495 AN cases, 2,688 OCD cases and 18,013 controls. We confirmed a high genetic correlation between AN and OCD (r_g_ = 0.49 ± 0.13, *p* = 9.07×10^−7^) and a sizable SNP heritability (SNP *h*^2^ = 0.21 ± 0.02) for the cross-disorder phenotype. Although no individual loci reached genome-wide significance, the cross-disorder phenotype showed strong positive genetic correlations with other psychiatric phenotypes (e.g., bipolar disorder, schizophrenia, neuroticism) and negative correlations with metabolic phenotypes (e.g., BMI, triglycerides). Follow-up analyses revealed that although AN and OCD overlap heavily in their shared risk with other psychiatric phenotypes, the relationship with metabolic and anthropometric traits is markedly stronger for AN than for OCD. We further tested whether shared genetic risk for AN/OCD was associated with particular tissue or cell-type gene expression patterns and found that the basal ganglia and medium spiny neurons were most enriched for AN/OCD risk, consistent with neurobiological findings for both disorders. Our results confirm and extend genetic epidemiological findings of shared risk between AN and OCD and suggest that larger GWASs are warranted.

## INTRODUCTION

Anorexia nervosa (AN) and obsessive-compulsive disorder (OCD) are heritable neuropsychiatric disorders with complex genetic etiologies.^1-3^ A wealth of genetic epidemiological findings supports associations between AN and OCD, suggesting the possibility of shared genetic risk factors. For example, ~20-50% of individuals with AN have a lifetime history of OCD^4-6^ and ~5-10% of patients with OCD have a lifetime history of AN,^7-10^ well above their population prevalences (OCD: ~2%,^11^ AN: ~1%^12^). In addition, first-degree relatives of AN probands have an approximately 3-5 fold elevated risk of OCD compared with relatives of healthy controls,^13-16^ further highlighting a potential genetic link between these disorders.

Recent genome-wide association studies (GWAS) of AN^17-29^ and OCD^20-22^ have provided important clues regarding the highly polygenic architecture of these two disorders. The largest AN GWAS published to date included 3,495 cases and 10,982 controls as part of the Eating Disorders Working Group of the Psychiatric Genomics Consortium (PGC-ED). In addition to the identification of the first genome-wide significant variant associated with AN risk, significant genetic correlations for AN with multiple psychiatric, neurocognitive, and metabolic phenotypes were also reported.^19^ Although there were no genome-wide significant variants identified in the OCD GWAS meta-analysis (2,688 cases and 7,037 controls), polygenic risk scores calculated in part of the sample significantly predicted OCD status in the remaining samples.^22^ A recent study using a subset of the GWAS data^17, 20^ reported a strong genetic correlation (*r_g_*) between AN and OCD (*r_g_* = 0.55, P = 2 ×10^−5^)^23^. This result mirrored the epidemiological findings summarized above as well as bivariate twin models that revealed a significant genetic overlap between selfreported OCD and AN (r_a_ = 0.52).^13^

Taken together, the available clinical, epidemiological and genetic results strongly point toward a shared etiology for AN and OCD, but our knowledge of the nature of this shared risk is limited. Thus, the aim of this study was to examine the cross-disorder genetic architecture of AN and OCD to gain a better understanding of their shared genetic etiology.

## MATERIALS AND METHODS

### Datasets

The genetic data for this study came from the summary statistics of the largest and most recently published GWAS for AN^19^ and OCD.^22^ There was no sample overlap between these studies according to SNP-based relatedness testing. For AN, we used data from a GWAS of 3,495 cases and 10,982 controls. For OCD, we used data derived from a similar recently published GWAS of 2,688 OCD cases and 7,031 controls. All participants were of European ancestry. Using these datasets, we conducted a meta-analysis of 6,183 AN-OCD cases and 18,013 controls. Although records regarding comorbidities are incomplete, and exist for approximately 50% of the OCD and 35% of the AN sample, we have identified 101 definite and 24 probable cases of AN among the OCD sample and 447 AN cases with self-reported OCD diagnoses. More information about these datasets can be found in the Supplementary Information.

### AN-OCD Cross-Disorder GWAS Meta-Analysis

As a part of the pre-processing step prior to meta-analysis, we identified all variants for which post-imputation dosage information was available to a sufficient degree and at an adequate level of quality at the call level in both datasets. To do this, we identified SNPs that met the following criteria for each dataset: (1) minor allele frequency (MAF) > 0.01; (2) imputation quality (INFO) score > 0.6; (3) ≥60% of cases and controls from the overall meta-analysis per trait available for each variant; and (4) not an A/T or G/C variant with frequencies between 0.4-0.6. A total of 90.0% (9,586,433 of 10,641,224) of the variants in the AN dataset and 91.6% (7,707,306 of 8,410,839) of the variants in the OCD dataset met these criteria. Following additional preprocessing of the summary statistics files (Supplementary Information), we identified 7,461,827 variants which were present in both datasets.

The meta-analysis used the Ricopili pipeline (https://sites.google.com/a/broadinstitute.org/ricopili), which is the standard analysis pipeline for the Psychiatric Genomics Consortium (PGC). This pipeline utilizes METAL^24^ and takes into account sample sizes as well as strength in direction of effect in each separate dataset (inverse-variance weighted meta-analysis using a fixed-effects model).

### AN and OCD heritability, intercept and genetic correlation estimates

We used linkage disequilibrium (LD) score regression^25^ to evaluate the relative contribution of polygenic effects and confounding factors (e.g., cryptic relatedness or population stratification) to deviation from the null in the genome-wide distribution of GWAS *χ^2^* statistics. Analysis was performed using pre-computed LD scores from European-ancestry samples in the HapMap phase 3 project. To estimate the impact of confounding factors, we evaluated the deviation of the LD score intercept from one, the expected value for a GWAS showing no sign of population stratification. LD score regression was also used to estimate SNP heritability.^25^ For AN, OCD, and AN-OCD cross-disorder phenotypes, we estimated liability-based heritability using population prevalence estimates of 0.9%, 2.5% and 3%, respectively. We also computed the genetic correlation between AN and OCD in order to determine whether the correlation is consistent with the previously reported high correlation between the two disorders.^23^

### Genetic Correlation of AN-OCD Single and Cross-Disorder Phenotypes with Other Traits

The summary statistics files from AN, OCD, and AN/OCD cross-disorder analyses were uploaded to LD Hub server (v1.4.1; http://ldsc.broadinstitute.org). For each summary statistic file, we computed pairwise genetic correlations with 231 datasets, each representing a particular GWAS phenotype.^26^ Subsequent exploratory analyses of the computed genetic correlations were then carried out, with the goal of describing consistent and unique pairwise correlations with other traits.

We first asked if there were traits that have significant correlations with the AN-OCD crossdisorder data at a multiple testing-corrected level of significance. We applied a stringent Bonferroni correction based on a total of 231 × 3 = 693 separate tests performed, with a *p*-value significance threshold of 7.2×10^−5^. We identified all single tests that passed this threshold, taking note of traits that did not meet the threshold in AN and OCD cohorts alone, but rather in the combined AN-OCD cohort. Second, we assessed the directionality of correlations between AN, OCD, and the tested traits, as well as identified traits where the correlations were in opposite directions. To analyze the concordance of correlation directions between AN and OCD, we constructed a sign test by stratifying traits into bins based on *p*-values (AN and OCD ≥ 0.05; AN or OCD < 0.05; AN or OCD < 1×10^−3^; AN or OCD < 1×10^−5^). For each bin, we counted the number of correlations which were in the same direction for AN and OCD, and binomial tests were performed under the null hypothesis that 50% of correlations should be in the same direction. We extracted traits for which: (1) AN or OCD had a correlation *p*-value that at least trended towards significance (*p* < 0.1); and (2) AN and OCD correlations had opposite signs. We hypothesized that the traits that came out of this exploration could potentially capture some of the genetic signal specific to each disorder rather than the cross-disorder shared risk.

Additionally, we performed tests to determine if there was evidence for sets of traits for which concordant correlations with AN and OCD were biased in magnitude towards one phenotype over the other. We identified 80 traits that had concordant direction of correlation between AN and OCD, as well as a correlation *p*-value < 0.05 in at least one of the two phenotypes. We binned these traits into categories as defined in LD Hub output and selected a subset of 10 categories for formal testing where we had at least three traits that fit the criteria above. For each category, we tested the null hypothesis that the number of traits where the magnitude of the correlation is greater in AN matches the number of traits where the magnitude is greater in OCD using a two-sided binomial test with an expected fraction of 0.5.

### Gene-based and pathway-based tests

We used MAGMA v1.06^27^ to conduct gene-based and pathway-based association tests on the AN-OCD cross-disorder genotype data. Association was tested using the SNP-wise mean model, in which the sum of −log(SNP *p*-value) for SNPs located within the transcribed region (defined using NCBI 37.3 gene definitions) was used as test statistic. Included variants had MAF > 0.01, and INFO scores > 0.8. All gene-based tests were performed on loci extending from 35kb upstream of transcription start site to 10kb downstream of the transcription end site. MAGMA accounts for gene-size, number of SNPs in a gene and LD between markers when estimating gene-based *p*-values. LD correction was based on estimates from the 1000 Genome phase 3 European ancestry samples. This yielded a total of 18,279 gene-based tests, with a multiple-test-corrected significance level of 0.05/18,279 = 2.74 × 10^−6^.

These gene-based p-values were used to analyze sets of genes in order to test for enrichment of association signals in genes belonging to specific biological pathways or processes. In the analysis, only genes on autosomes, and genes located outside the broad major histocompatibility complex (MHC) region (hg19:chr6:25-35Mb) were included. For gene sets we used sets with 10-1000 genes from the Gene Ontology sets curated from MsigDB 6.0,^28, 29^ yielding a total of 17,081 separate pathway-based tests, with a multiple-test-corrected significance threshold of 0. 05/17,081 = 2.92 × 10^−6^.

### Partitioned LD score regression

Heritability was partitioned into different genomic regions using LD score regression.^25, 30^ LD score regression can estimate common SNPs heritability by regressing the *χ^2^* association statistics against the LD scores, which are defined as the sum of ***r^2^*** for each SNP. Indeed, assuming polygenicity, SNPs in high LD should on average be more associated with the trait than SNPs in low LD because they are expected to be in linkage with more causal variants. The relationship between *χ^2^* statistics and LD scores is directly dependent on the proportion of genetic variance of the trait, which allows to estimate heritability.

Partitioned LD score regression uses the same principle except that SNPs are partitioned into diverse functional categories. Indeed, if some categories are enriched in causal variants, the relationship between *χ^2^* statistics and LD scores should be stronger than for categories with few causal variants, allowing to estimate heritability enrichment of diverse functional categories. We used the “baseline model” provided on the partitioned LD score regression website (https://github.com/bulik/ldsc/wiki/Partitioned-Heritability) to compute heritability enrichment for 28 functional categories.

### Tissue and cell type association

We used MAGMA^27^ and partitioned LD score regression^30^ to look for association between tissue/cell type specificity in gene expression and gene-level genetic association with AN-OCD. The methods for these analyses were extensively described elsewhere.^31^ Briefly, for tissue level expression, we downloaded the median RPKM across individuals (V6p) from the GTEx portal (https://www.gtexportal.org). For the single cell RNA-seq, we examined single cell RNA-seq data from 9,970 single cells coming from five different mouse brain regions (cortex, hippocampus, striatum, midbrain and hypothalamus) allowing the identification of 24 different brain cell types. The prototypical expression of each cell type was obtained by averaging gene expression levels for all cells classified in each of the 24 cell types. We then obtained a measure of specificity for each gene and each cell type/tissue by dividing the expression of each gene in each cell type/tissue by the total expression of the gene.

For MAGMA, we binned the specificity measure for each cell type into 41 bins (0 for genes not expressed, 1 for genes below the 2.5% quantile in specificity,…, 40 for genes between the 97.5% quantile in specificity and the 100% quantile). We then tested for a positive relationship between the specificity bins in each cell type/tissue and the gene-level genetic association of each gene (-10kb upstream to 1.5kb downstream). For LD score regression, we binned the specificity measure into 11 bins for each cell type (0 for genes not expressed, 10 for genes between the 90% quantile and the 100% quantile) and tested whether genomic regions around genes (-10kb downstream to +1.5 kb upstream) in the top 10% most specific genes for each cell type/tissue were enriched for AN-OCD cross-disorder heritability.

## RESULTS

### Cross-Disorder Meta-Analysis

The AN-OCD meta-analysis included 6,183 cases, 18,031 controls, and 7,461,827 variants. As shown in the quantile-quantile (QQ) plot (***Figure 1a***), the genomic inflation factor (*λ*_1000_) of 1.010 indicates no signs of excessive inflation. Inflation statistics from individual AN^19^ and OCD^22^ datasets appear similarly well controlled (*λ*_1000_ values of 1.011 and 1.009 respectively). As shown in the Manhattan plot in ***Figure 1b***, no variants reached genome-wide significance and 30 loci had a lead SNP with *p* < 10^−5^. The most significant peaks were located near genes *CXCR4* (chr2), *KIT* (chr4), the MHC (chr6), *FAM19A2* (chr12), and *SLC25A42* (chr19). The most prominent signal that was driven by both AN and OCD lies near the MHC (index SNP rs75063949, chr6:25591041 in *LRRC16A* (***Figure S1***). As shown in the forest plot for this SNP (***Figure S1a)***, the 95% confidence interval on the natural log of the odds ratio excludes 0 in both AN and OCD. Another cross-disorder signal was found upstream of *KIT* on chr4 (***Figure S2***), while the locus found within *FAM19A2* on chr12 appears primarily driven by AN (***Figure S3***).

**Figure 1.**
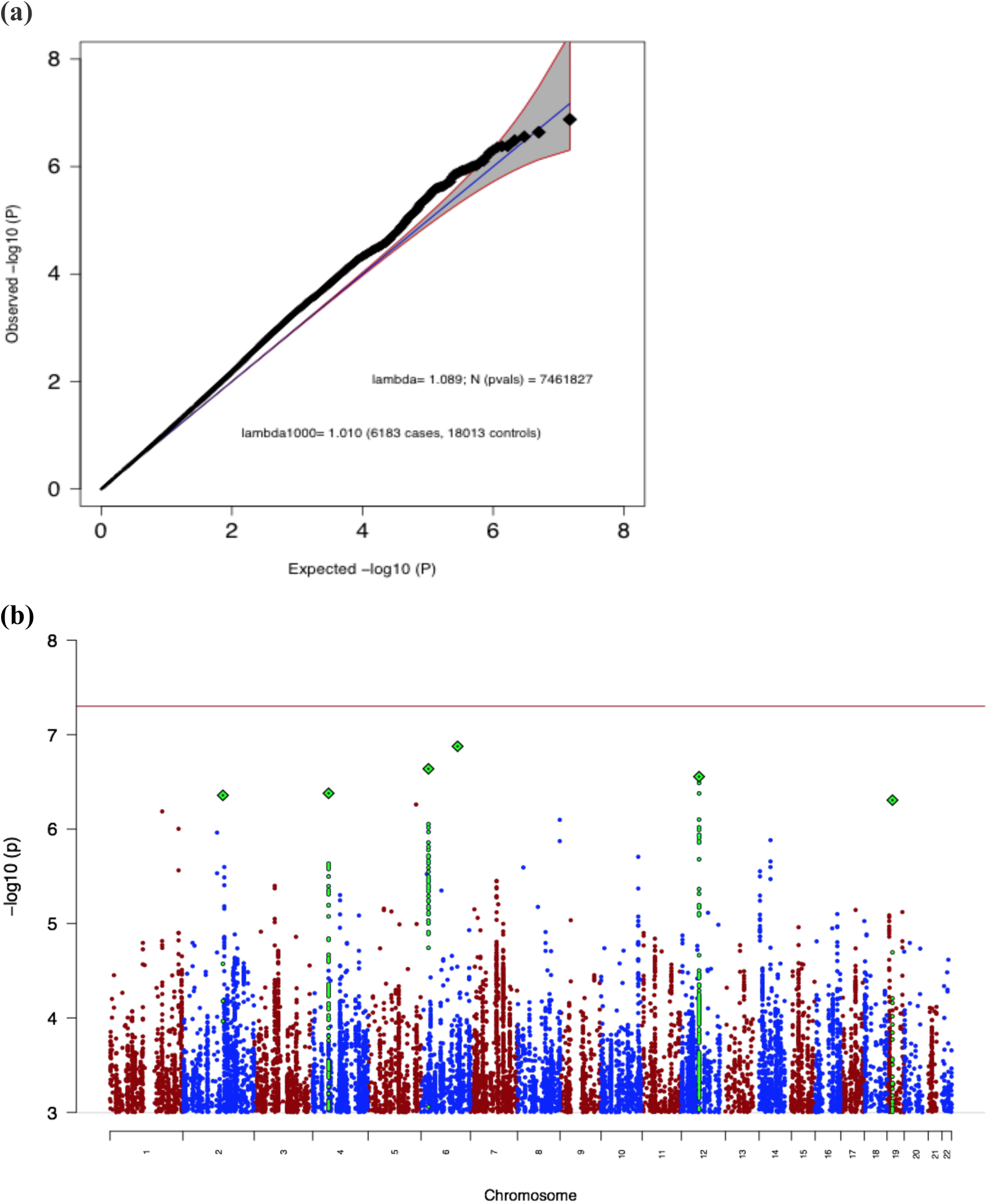
**(a)** Quantile-quantile (QQ) plot of the AN-OCD cross-disorder meta-analysis, showing a slight departure from a null model of no associations (blue line and grey 95% confidence interval). **(b)** Manhattan plot of the AN-OCD cross-disorder meta-analysis of 6,183 cases and 18,031 controls. The x-axis shows genomic position (chr1-chr22), and the y-axis shows statistical significance as −log10(P). The red line shows the genome-wide significance threshold (P=5×10^−8^).

### Heritability and Genetic Correlation Estimates for AN and OCD

As an additional assessment of the distribution of test statistics for our constructed cross-disorder meta-analysis, we computed liability-based heritability estimates for AN and OCD datasets, as well as the genetic correlation between the two disorders based on the disorder-specific summary statistics. Using population prevalence estimates of 0.9%, 2.5%, and 3% for AN, OCD and the AN-OCD cross-disorder phenotype respectively, we computed heritability estimates of 0.18 for AN, 0.29 for OCD, and 0.21 for AN-OCD (***Table 1***). The AN and OCD estimates were computed only using the SNPs present in both datasets and are consistent with previously reported estimates of 0.20 for AN^19^ and 0.28 for OCD.^22^ The LD Score regression intercept of the cross-disorder association analysis was 1.01, therefore providing further evidence against the presence of genomic inflation.

**Table 1.** LD Score regression results. Shown are the SNP-based heritabilities for AN, OCD, and AN-OCD cross-disorder phenotype (rounded to two decimal points).

|  | **AN** | **OCD** | **AN-OCD** |
| --- | --- | --- | --- |
| **Liability Scale $h^2$** | 0.18 | 0.29 | 0.21 |
| **Standard Error for $h^2$** | 0.03 | 0.04 | 0.02 |
| **Mean Chi-Square** | 1.08 | 1.06 | 1.11 |
| **LD Score Regression Intercept** | 1.01 | 0.99 | 1.01 |

Using the largest published datasets to date, we computed a genetic correlation between AN and OCD of 0.49 (*se* = 0.13; *p* = 9.07×10^−7^) and therefore supporting the previously reported genetic correlation calculated using a subset of these samples.^23^

### Genetic Correlations with Other Phenotypes

Following Bonferroni correction, 19 of the 231 phenotypes available in LD Hub were significantly correlated with AN-OCD shared genetic risk. Among psychiatric phenotypes, bipolar disorder, schizophrenia, and PGC cross-disorder shared risk (comprising bipolar disorder, schizophrenia, autism spectrum disorders, attention deficit/hyperactivity disorder, and major depressive disorder) were positively correlated with AN-OCD shared risk (***Figure 2***). These findings are in line with previous work,^23^ including the most recent AN GWAS^19^ and emphasize the shared genetic architecture of psychiatric disorders in general. We also observed a positive correlation between AN-OCD cross-disorder risk and years of schooling, which mirrors the positive genetic correlation reported for AN and educational attainment.^19^

**Figure 2.**
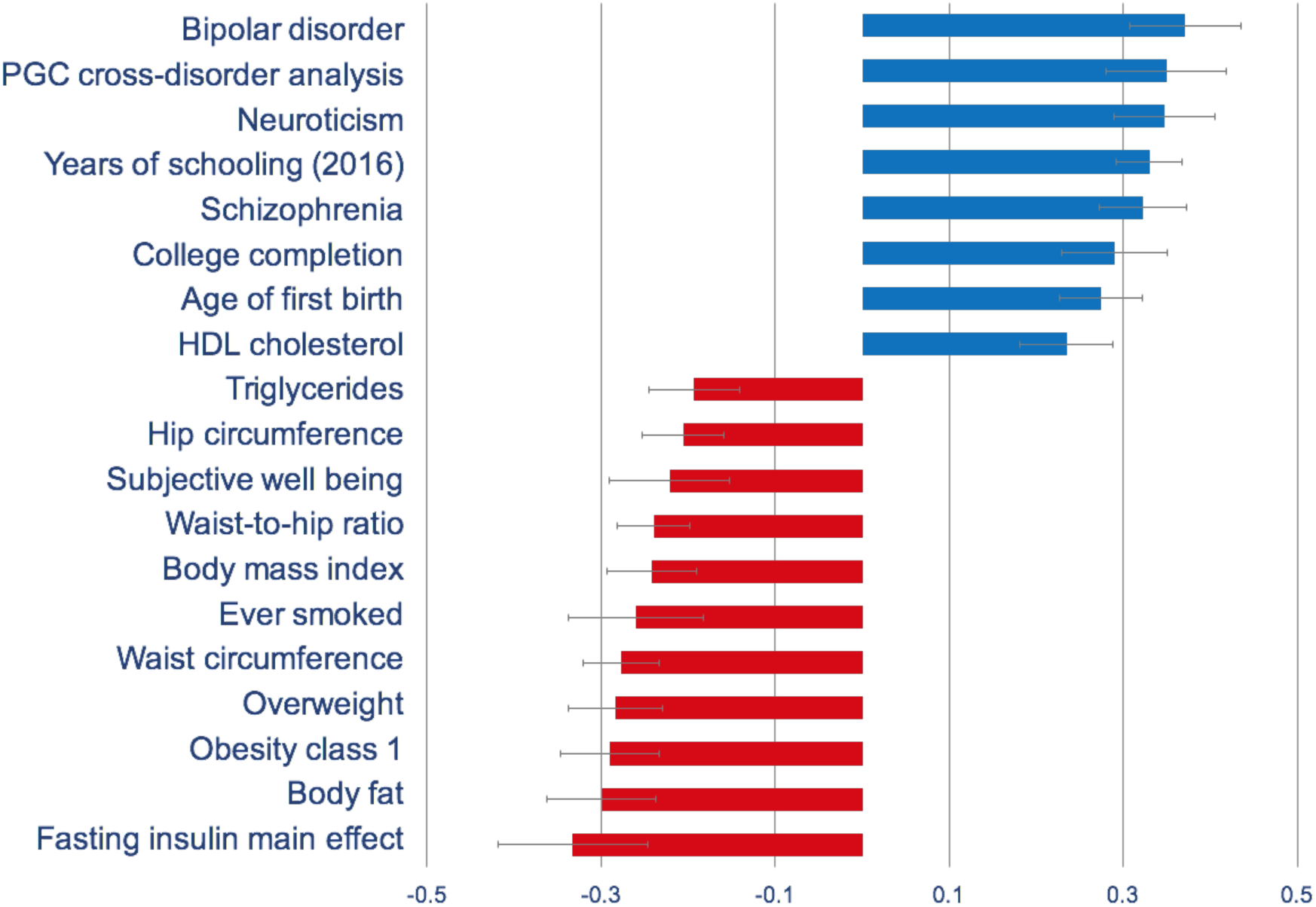
Genetic correlations of the AN-OCD cross-disorder phenotype with psychiatric, metabolic, and other phenotypes. Only significant results after Bonferroni correction are shown

For metabolic traits, high body mass index and related phenotypes (i.e., class I obesity, overweight, hip circumference, waist-to-hip ratio) were negatively correlated with AN-OCD shared risk. Similarly, whereas unfavorable metabolic markers such as triglyceride and fasting insulin yielded negative genetic correlations with AN-OCD, HDL cholesterol (which is a favorable metabolic marker) yielded a significant positive genetic correlation. These findings also support the genetic correlations previously reported for AN,^19^ therefore suggesting a metabolic component in the genetic etiology of AN as well as AN-OCD cross-disorder phenotype.

Many of the stronger, higher confidence pairwise correlations derived from LD Hub appear to be shared by AN and OCD. In visualizing the correlations, we first noted this trend, in that there appeared to be very few correlations of particularly high confidence where the directions of correlation pointed in opposite directions between the two disorders (***Figure 3a***). To formally test this, we assessed the frequency of the same direction correlations across different p-value thresholds, with an expected value of 50% under the null, and found that more significant results were more concordant, as expected (***Figure 3b***). This observation is consistent with the observed high genetic correlation between AN and OCD.

**Figure 3.**
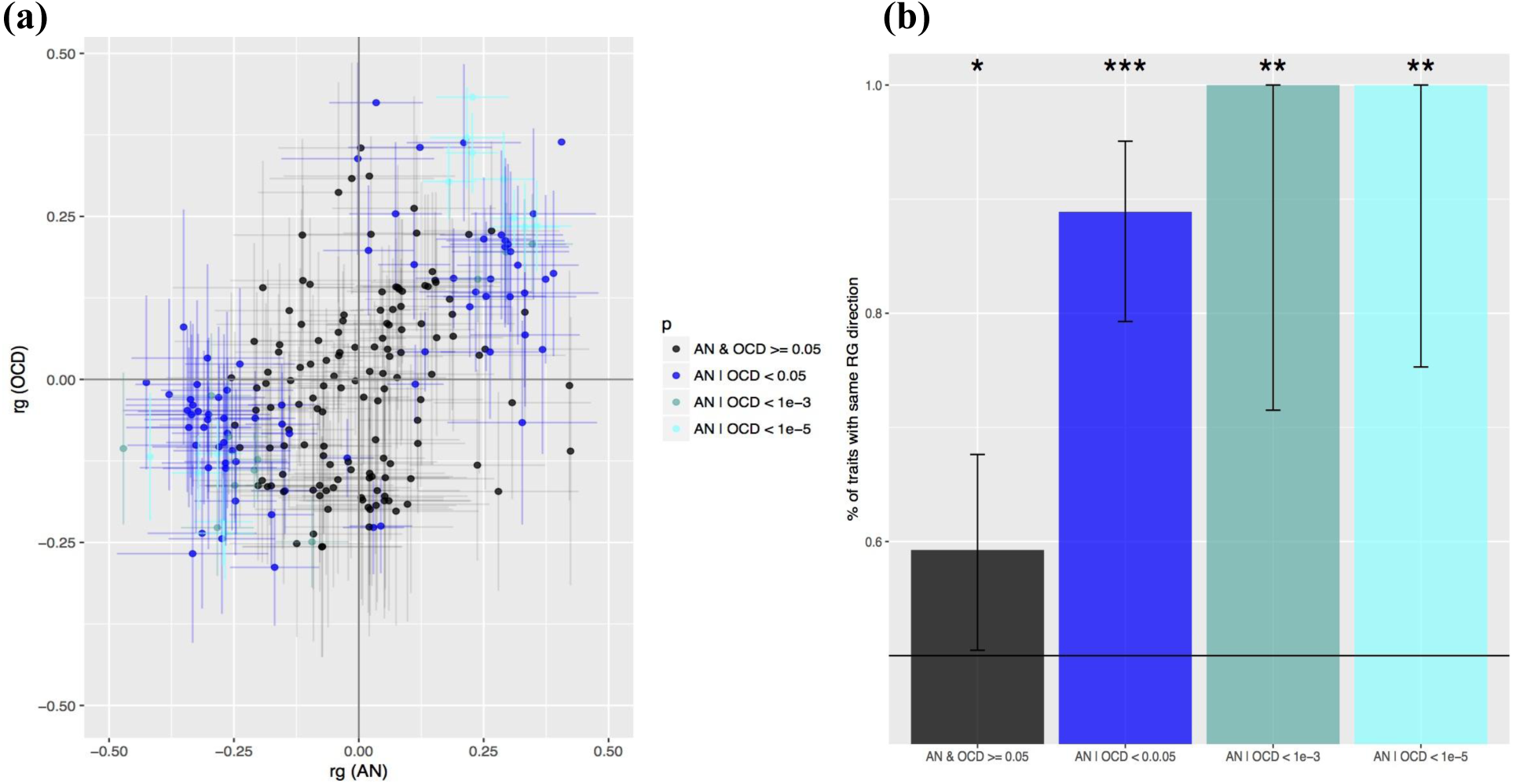
AN and OCD show overlapping patterns of genetic correlation across 229 traits. **(a)** Distribution of pairwise correlations of traits with AN and OCD in two-dimensional space. Bars depict one standard error in each direction for the pairwise correlation estimate between each trait in question and AN or OCD, with lines stretched horizontally in reference to AN and vertically in reference to the OCD estimate. Data points are color coded based on the p-value bin they fall into, listed in the legend. **(b)** Results of correlation sign test between AN and OCD values, for traits previously binned and color coded. Bar represent the 95% confidence interval. For indicators of significance, “***” indicates p < 0.0001, “**” indicates p < 0.001, and “*” indicates p < 0.05.

In order to determine if there were traits with *p* < 0.1 where the correlations with AN and OCD were markedly different, we identified 16 traits where the signs of the genetic correlations with AN and with OCD were opposite (***Figure S4***). Many traits were metabolic and anthropometric in nature, for which the correlations with AN were much stronger. There were two phenotypes for which the correlations with AN and OCD were both notably high but in opposite directions. These included the personality trait “openness to experience” (correlation positive with AN but negative with OCD), and genetic risk for Amyotrophic Lateral Sclerosis (also positive in AN but negative in OCD).

Given that many of the stronger correlations were in the same direction, we then examined whether particular phenotypes—clustered by some higher-order category classification—would be enriched for stronger correlation magnitudes in one phenotype versus the other. To test this, we took the subset of traits where the direction of correlation with AN and with OCD were concordant, and at least one phenotype between AN and OCD had a nominally significant correlation at *p* < 0.05. We next took the higher-order category classifications in LD Hub that had been applied to each trait, and took the subset of categories where we had three or more applicable traits that met the previously described criteria (10 categories total). For each category we tested the null hypothesis that 50% of traits in that category would have a greater correlation magnitude in OCD, again using a two-sided binomial test. We found that correlation magnitudes were heavily shifted towards AN for anthropometric and metabolic categories (***Figure S5***). Consistent with this observation, lipid and glycemic categories show similar bias towards AN, albeit with less statistical power due to a smaller number of traits represented in these categories.

### Partitioned heritability

We also investigated whether particular genomic annotations were enriched in AN-OCD crossdisorder heritability using partitioned LD score regression.^30^ As expected with the current sample size, we did not observed any significant enrichments at a 5% false discovery rate (***Figure S6***), and the most significant annotation was conservation regions across 29 mammals^32^ with an estimated 17-fold heritability enrichment (*q*-value = 0.23). Since this annotation was previously shown to be enriched in heritability for a variety of traits,^30^ it is likely that the lack of significant association in our study is due to limited statistical power.

### Tissue and cell-type association

We next aimed to detect tissues involved in the aetiology of AN-OCD using MAGMA.^27^ We did not observe any tissue types in GTEx^33^ significantly associated with AN-OCD following Bonferroni correction; however, the strongest associations were observed with brain tissues. Notably, the top hit in GTEx was the nucleus accumbens from the basal ganglia (***Figure S7***).

Given the suggestive evidence of the implication of brain tissues in the etiology of AN-OCD in our previous analysis, we used MAGMA to investigate whether there was an association between AN-OCD cross-disorder risk and any of 24 mouse brain cell types.^31^ We observed that medium spiny neurons (MSNs) were significantly associated with AN-OCD after Bonferroni correction (***Figure 4***). As MSNs are located in the basal ganglia, the association of MSNs with AN-OCD is concordant with the tissue-level association of the basal ganglia using GTEx data. Of note, we did not observe the same significant association when we used LD score regression (***Figure S8***), indicating that this association may not be very robust (which is not surprising given the current sample size).

**Figure 4.**
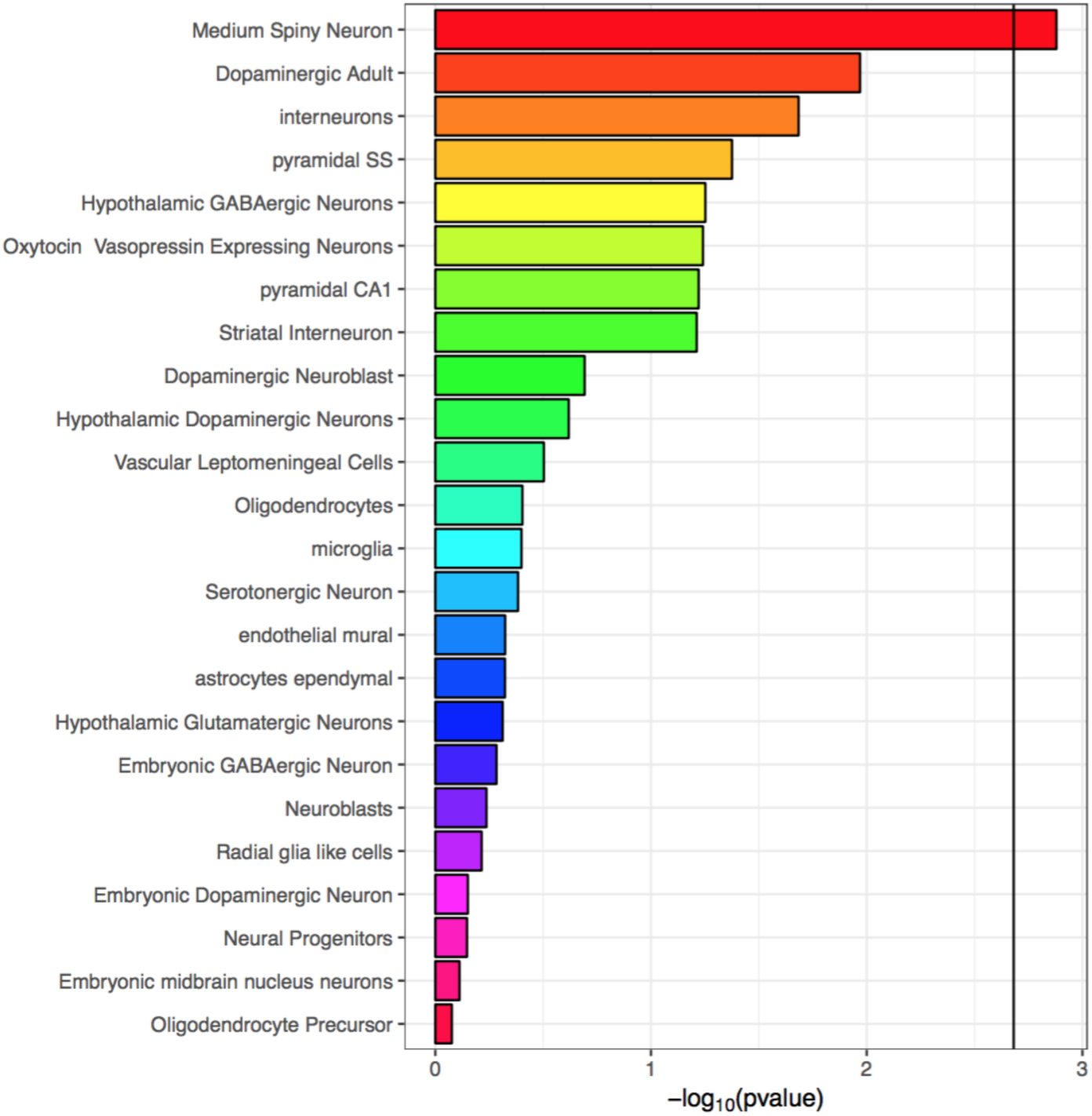
Association between mouse brain cell-type specific expression and gene-level genetic association to AN/OCD using MAGMA. The black vertical bar represents the Bonferroni significant threshold.

### Additional analyses

We performed gene-based and pathway-based tests on the AN/OCD cross disorder summary statistics using the MAGMA package (see methods for details). One gene (*KIT*) passed the multiple test correction threshold of 2.5 × 10^−6^ (***Figure S9***). We did not detect any pathways that survived multiple test correction for 17,081 separate pathway-based tests.

## DISCUSSION

This cross-disorder GWAS of AN and OCD was geared toward explicating the shared genetic architecture of the two disorders. The clinical and epidemiological connections between AN and OCD have been well-documented,^4-16^ and it is especially important to examine the potential genetic connection between these two disorders in the light of recent genetic and twin-based findings.^13^ As disorder-specific association studies have been conducted (and analyses with larger sample sizes are ongoing), the present study was geared toward examining the shared AN-OCD risks as opposed to the dissection of any disorder-specific risk factors.

The combined GWAS meta-analysis of 6,183 cases and 18,013 controls did not yield any genome-wide significant findings for shared AN-OCD risk. Although there is one genome-wide significant risk locus identified for AN, the available OCD and AN samples continue to be small relative to those needed for the reliable detection of genome-wide significant loci. Therefore, this AN-OCD analysis was likely underpowered to detect genome-wide significant findings and genetic heterogeneity between AN and OCD may also play a role. Thus, in the absence of significant hits, we describe the loci for which we found strongest evidence for association. The most prominent reliable signal lies near the edge of the MHC in the gene *LRRC16A* (***Figure S1a***) and is driven by both AN (p = 4.19 × 10^−5^) and OCD (p = 1.53 × 10^−3^). Although this gene has no clear connection to brain phenotypes, and it remains to be seen whether this signal will reach significance, it is notable that a role for autoimmunity has previously been implicated in both AN^34^ and OCD.^35^

Another association driven by both AN (*p* = 1.62 × 10^−6^) and OCD (*p* = 0.011) is located just upstream of *KIT*, which is thought to be involved in cell growth and differentiation.^36^ We were unable to find literature supporting a link between this gene and a psychiatric phenotype. Gene-based association for *KIT* barely exceeded the pre-determined significance threshold, though the regional association plots suggest a true signal and not one that is artefactual, so more data are needed before seriously considering its role in AN/OCD. Taken together, our results strongly point to the possibility of identifying loci for shared AN/OCD risk as sample sizes for both disorders increase.

We observed a genetic correlation of 0.49 (*se* = 0.13; *p* = 9.07×10^−7^) between AN and OCD, supporting previous results^23^ computed using smaller AN and OCD datasets. We also examined the genetic correlations of AN and OCD with a wide range of other phenotypes. The results of these analyses hinted at some of the similarities and potential differences between AN and OCD genetic architecture. Although it is clear that both share correlations with other psychiatric disorders and that many of the particularly strong correlations with the traits explored are common between the two disorders, we note a markedly stronger correlation in AN with metabolic and anthropometric traits. It is possible that a subset of these correlations may represent a component of AN genetic architecture, which may distinguish AN from OCD and set AN apart from other psychiatric disorders as having both a psychiatric and a metabolic etiology.^19^ As previously noted, only greatly expanded GWAS studies of both AN and OCD will be able to tell us what genetic risk factors are distinct to AN and to OCD.

Tissue and cell type analyses suggested that the basal ganglia and MSNs play a role in AN-OCD cross-disorder risk. Larger samples, however, are needed to conclusively implicate specific brain regions and cell types as our findings were not robust enough when using two different analytical methods. Nevertheless, the association of MSNs seems plausible as these neurons have also been associated with schizophrenia^31^ and neuroticism,^37^ both of which were found to be significantly genetically correlated with AN-OCD shared risk in our analyses. A large body of literature supports a role for the basal ganglia and its associated substructures in both AN and OCD. In AN, the nucleus accumbens is involved in feeding behavior,^38^ hyperactivity^39^ and its ablation increased drive to eat, in some individuals with AN.^40^ In OCD, neuroimaging studies have repeatedly shown hyperactivity in the head of the caudate nuclei^41^ and deep brain stimulation targeting striatal areas is often therapeutic,^42^ both consistent with the cortico–striato–thalamo–cortical (CSTC) model of OCD.^43^

The strengths of this study include the use of the largest and most current GWAS results for both AN and OCD and rigorous pre-processing of the summary statistics files prior to analyses, to ensure comparability in this first ever AN-OCD cross disorder GWAS. Limitations of this study include heterogeneity, the fact that both the AN and OCD GWAS are currently underpowered and our data on comorbidities are limited. Future directions for this work as sample size increases will include the incorporation of subphenotype information in the analyses as well as Mendelian randomization analyses to examine causal inference.

## ACKNOWLEDGEMENTS

Statistical analyses were carried out on the NL Genetic Cluster Computer (http://www.geneticcluster.org) hosted by SURFsara.

## CONSORTIA

### Eating Disorders Working Group of the PGC

Roger Adan, Tetsuya Ando, Jessica Baker, Andrew Bergen, Wade Berrettini, Andreas Birgegård, Claudette Boni, Vesna Boraska Perica, Harry Brandt, Roland Burghardt, Matteo Cassina, Carolyn Cesta,Maurizio Clementi, Jonathan Coleman, Roger Cone, Philippe Courtet, Steven Crawford, Scott Crow, James Crowley, Unna Danner, Oliver Davis, Martina de Zwaan, George Dedoussis, Daniela Degortes, Janiece DeSocio, Danielle Dick, Dimitris Dikeos, Monika Dmitrzak-Weglarz, Elisa Docampo, Karin Egberts, Stefan Ehrlich, Geòrgia Escaramís, Tõnu Esko, Xavier Estivill, Angela Favaro, Fernando Fernández-Aranda, Manfred Fichter,Chris Finan, Krista Fischer, Manuel Föcker, Lenka Foretova, Monica Forzan, Christopher Franklin, Héléna Gaspar, Fragiskos Gonidakis, Philip Gorwood, Monica Gratacos, Sébastien Guillaume, Yiran Guo, Hakon Hakonarson, Katherine Halmi, Konstantinos Hatzikotoulas, Joanna Hauser, Johannes Hebebrand, Sietske Helder, Judith Hendriks, Beate Herpertz-Dahlmann, Wolfgang Herzog, Christopher Hilliard, Anke Hinney, Laura Huckins, James Hudson, Julia Huemer, Hartmut Imgart, Hidetoshi Inoko, Susana Jiménez-Murcia, Craig Johnson, Jenny Jordan, Anders Juréus, Gursharan Kalsi, Debora Kaminska, Allan Kaplan, Jaakko Kaprio, Leila Karhunen, Andreas Karwautz, Martien Kas, Walter Kaye, James Kennedy, Martin Kennedy, Anna Keski-Rahkonen, Kirsty Kiezebrink, Youl-Ri Kim, Kelly Klump, Gun Peggy Knudsen, Bobby Koeleman, Doris Koubek, Maria La Via, Mikael Landén, Robert Levitan, Dong Li, Paul Lichtenstein, Lisa Lilenfeld, Jolanta Lissowska, Pierre Magistretti, Mario Maj, Katrin Mannik, Nicholas Martin, Sara McDevitt, Peter McGuffin, Elisabeth Merl, Andres Metspalu, Ingrid Meulenbelt, Nadia Micali, James Mitchell, Karen Mitchell, Palmiero Monteleone, Alessio Maria Monteleone, Grant Montgomery, Preben Mortensen, Melissa Munn-Chernoff, Benedetta Nacmias, Ida Nilsson, Claes Norring, Ioanna Ntalla, Julie O’Toole, Jacques Pantel, Hana Papezova, Richard Parker, Raquel Rabionet, Anu Raevuori, Andrzej Rajewski, Nicolas Ramoz, N. William Rayner, Ted Reichborn-Kjennerud, Valdo Ricca, Stephan Ripke, Franziska Ritschel, Marion Roberts, Alessandro Rotondo, Filip Rybakowski, Paolo Santonastaso, André Scherag, Ulrike Schmidt, Nicholas Schork, Alexandra Schosser, Jochen Seitz, Lenka Slachtova, P. Eline Slagboom, Margarita Slof-Op’t Landt, Agnieszka Slopien, Tosha Smith, Sandro Sorbi, Eric Strengman, Michael Strober, Patrick Sullivan, Jin Szatkiewicz, Neonila Szeszenia-Dabrowska, Ioanna Tachmazidou, Elena Tenconi, Laura Thornton, Alfonso Tortorella, Federica Tozzi, Janet Treasure, Artemis Tsitsika, Konstantinos Tziouvas, Annemarie van Elburg, Eric van Furth, Tracey Wade, Gudrun Wagner, Esther Walton, Hunna Watson, D. Blake Woodside, Shuyang Yao, Zeynep Yilmaz, Eleftheria Zeggini, Stephanie Zerwas, Stephan Zipfel, Lars Alfredsson, Ole Andreassen, Harald Aschauer, Jeffrey Barrett, Vladimir Bencko, Laura Carlberg, Sven Cichon, Sarah Cohen-Woods, Christian Dina, Bo Ding, Thomas Espeseth, James Floyd, Steven Gallinger, Giovanni Gambaro, Ina Giegling, Stefan Herms, Vladimir Janout, Antonio Julià, Lars Klareskog, Stephanie Le Hellard, Marion Leboyer,Astri Lundervold, Sara Marsal, Morten Mattingsdal, Marie Navratilova, Roel Ophoff, Aarno Palotie, Dalila Pinto, Samuli Ripatti, Dan Rujescu, Stephen Scherer, Laura Scott, Robert Sladek, Nicole Soranzo, Lorraine Southam, Vidar Steen, H-Erich Wichmann, Elisabeth Widen, Co-Chairs: Gerome Breen, Cynthia Bulik

### Obsessive Compulsive Disorder Working Group of the PGC

Paul D Arnold, Kathleen D Askland, Cristina Barlassina, Laura Bellodi, O J Bienvenu, Donald Black, Michael Bloch, Helena Brentani, Christie L Burton, Beatriz Camarena, Carolina Cappi, Danielle Cath, Maria Cavallini, David Conti, Edwin Cook, Vladimir Coric, Bernadette A Cullen, Danielle Cusi, Lea K Davis, Richard Delorme, Damiaan Denys, Eske Derks, Valsamma Eapen, Christopher Edlund, Lauren Erdman, Peter Falkai, Martijn Figee, Abigail J Fyer, Daniel A Geller, Fernando S Goes, Hans Grabe, Marcos A Grados, Benjamin D Greenberg, Edna Grünblatt, Wei Guo, Gregory L Hanna, Sian Hemmings, Ana G Hounie, Michael Jenicke, Clare Keenan, James Kennedy, Ekaterina A Khramtsova, Anuar Konkashbaev, James A Knowles, Janice Krasnow, Cristophe Lange, Nuria Lanzagorta, Marion Leboyer, Leonhard Lennertz, Bingbin Li, K-Y Liang, Christine Lochner, Fabio Macciardi, Brion Maher, Wolfgang Maier, Maurizio Marconi, Carol A Mathews, Manuel Matthesien, James T McCracken, Nicole C McLaughlin, Euripedes C Miguel, Rainald Moessner, Dennis L Murphy, Benjamin Neale, Gerald Nestadt, Paul Nestadt, Humberto Nicolini, Ericka Nurmi, Lisa Osiecki, David L Pauls, John Piacentini, Danielle Posthuma, Ann E Pulver, H-D Qin, Steven A Rasmussen, Scott Rauch, Margaret A Richter, Mark A Riddle, Stephan Ripke, Stephan Ruhrmann, Aline S Sampaio, Jack F Samuels, Jeremiah M Scharf, Yin Yao Shugart, Jan Smit, Barbara Stranger, Daniel Stein, S Evelyn Stewart, Maurizio Turiel, Homero Vallada, Jeremy Veenstra-VanderWeele, Michael Wagner, Susanne Walitza, Y Wang, Jens Wendland, Nienke Vulink, Dongmei Yu & Gwyneth Zai

